# Composition of estuarine sediment microbiome from a chronosequence of restored urban salt marshes

**DOI:** 10.1101/641274

**Authors:** Nathan Morris, Mary Alldred, Chester Zarnoch, Elizabeth Alter

## Abstract

Salt marshes play an important role in the global nutrient cycle. The sediments in these systems harbor diverse and complex bacterial communities possessing metabolic capacities that provide ecosystem services such as nutrient cycling and removal. On the East Coast of the United States, salt marshes have been experiencing degradation due to anthropogenic stressors. Salt marsh islands within Jamaica Bay, New York City (USA), are surrounded by a large highly urbanized watershed and have declined in area. Restoration efforts have been enacted to reduce further loss, but little is known about how microbial communities develop following restoration activities, or how processes such as nitrogen cycling are impacted. Sediment samples were collected at two sampling depths from five salt marsh islands to characterize the bacterial communities found in marsh sediment including a post-restoration chronosequence of 3-12 years. We used 16s rRNA amplicon sequencing to define alpha and beta diversity, taxonomic composition, and predicted metabolic profile of each sediment sample. We found significant differences in alpha diversity between sampling depths, and significant differences in beta diversity, taxonomic composition, and predicted metabolic capacity among the five sampling locations. The youngest restored site and the degraded natural sampling site exhibited the most distinct communities among the five sites. Our findings suggest that while the salt marsh islands are located in close proximity to each other, they harbor distinct bacterial communities that can be correlated with the post-restoration age, marsh health, and other environmental factors such as availability of organic carbon.

**IMPORTANCE:** Salt marshes play a critical role in the global nutrient cycle due to sediment bacteria and their metabolic capacities. Many East Coast salt marshes have experienced significant degradation over recent decades, thought largely to be due to anthropogenic stressors such as nitrogen loading, urban development, and sea-level rise. Salt marsh islands in Jamaica Bay (Queens/Brooklyn NY) are exposed to high water column nitrogen due to wastewater effluent. Several receding marsh islands have been subjected to restoration efforts to mitigate this loss. Little is known about the effect marsh restoration has on bacterial communities, their metabolic capacity, or how they develop post-restoration. Here we describe the bacterial communities found in marsh islands including a post-restoration chronosequence of 3-12 years and one degraded marsh island that remains unrestored.

## INTRODUCTION

Microbial communities drive essential ecosystem services in estuaries and salt marshes, most significantly via nutrient recycling and removal. For example, microbes carry out denitrification and anammox pathways, using nitrate (NO_3_^-^) or nitrite (NO_2_^-^) to produce di-nitrogen gas (N_2_) (1–4). These nitrogen (N) removal processes are particularly important in urban estuaries, which frequently suffer from eutrophication due to runoff and incomplete treatment of sewage (5). Conversely, N recycling can sustain eutrophic conditions through mineralization of sediment organic matter as well as dissimilatory nitrate reduction to ammonium (DNRA), a microbial metabolic pathway that reduces nitrate and nitrite to ammonium (NH_4_^+^). Denitrification is an important nitrogen pathway in salt marshes (6), but eutrophic environments may alter sediment microbial communities to favor N recycling over N removal (7–9).

The loss and degradation of salt marshes in estuaries has resulted in long-term restoration efforts to supplement or replace sediments and vegetation. A major goal of these restoration efforts is to maximize ecosystem services such as N retention and removal provided by microbial communities (10, 11). Despite this critically important objective, relatively little is known about the diversity of microbes in restored marshes, or the extent to which different clades are responsible for the metabolic processes involved in N cycling. Studies performed on natural marshes have shown that eutrophic conditions can alter microbial community structure and function (12–14). Other studies have documented the genomic potential of salt marsh microbes involved in sediment N cycling. Studies performed on salt marsh chronosequences have also demonstrated changes in microbial community composition (15) and N cycling processes (16, 17); however these studies were performed on a marsh chronosequence that had developed naturally over time, not due to restoration efforts. We know less about microbial community composition and diversity in restored estuarine sediments within urban ecosystems, and how these communities may change with increased age of the restored habitat. Constructed marshes may differ from natural marshes in microbial community composition due to the lack of accumulated organic carbon, fixed N, and differences in succession timelines (18). Urban constructed marshes are of particular interest because of their role in N removal in these highly eutrophic systems.

Jamaica Bay is an urban estuary surrounded by Brooklyn, Queens, and part of Nassau County (NY). The watershed of Jamaica Bay is ∼37,000 ha, and almost all of this area is highly urbanized (19). Approximately 15,785 kg of total dissolved nitrogen is exported to Jamaica Bay every day, primarily derived from wastewater treatment plant effluent (19). Current estimates of salt-marsh loss in Jamaica Bay are roughly 13 ha y^-1^ (20, 21), and most of this loss is from inland marsh islands, which have shrunk from 950 ha in 1951 (22) to 344 ha in 2013 (23). In an attempt to combat this loss, government entities including the National Park Service (NPS) and Army Corps of Engineers began marsh restoration projects in 2003 (23, 24). Since these projects have been carried out in different locations every few years (from 2003 through 2012), a chronosequence of restored salt marsh islands now exists in Jamaica Bay. In addition, because Jamaica Bay is well mixed these locations experience similar water-column chemistry providing an ideal system to investigate how microbial communities may change with age within eutrophic estuaries.

Management plans and ecosystem models for eutrophic estuaries require a better understanding of the composition and metabolic capacity of sediment microbial communities within restored marshes and how they change during marsh development. Here, we investigate the diversity of microbial communities found in sediments across the chronosequence of restored salt marsh islands in Jamaica Bay. Sediment was collected from five salt marsh islands in 2015, including four sites that at time of sampling were 3-12 years post-restoration (Fig. 1). Yellow Bar Marsh (YB), Elders West Marsh (EW), Elders East Marsh (EE), and Big Egg Marsh (BE) represent 3, 5, 9 and 12 years post-restoration, respectively. A fifth site, Black Bank Marsh (BB), > 200 years old (25, 26), has not yet been restored and is considered to be a degrading marsh due to erosion at the marsh edge, loss of vegetative cover, and reduction of belowground biomass (21, 23). At each site, we assessed vegetation and sediment characteristics, and collected sediment cores from 6 plots at two sampling depths (0-5 cm and 5-10 cm) for microbial community analysis. We used high-throughput sequencing of the 16S rRNA gene to describe and compare the diversity, taxonomic composition, and predicted functional capacity of these communities at two sediment depths. We predicted that: 1) microbial communities would differ between sampling depths and among sites along the chronosequence; 2) younger salt marshes would harbor more taxa related to autotrophic N fixation; and 3) as plant biomass and sediment organic content increased the microbial communities would shift to taxa with the metabolic capacity to break down organic material and remove N. Bacterial assemblages found in marsh sediments in various stages of development and at sites affected by intense anthropogenic pressure, should be indicative of marsh age.

**FIG 1.**
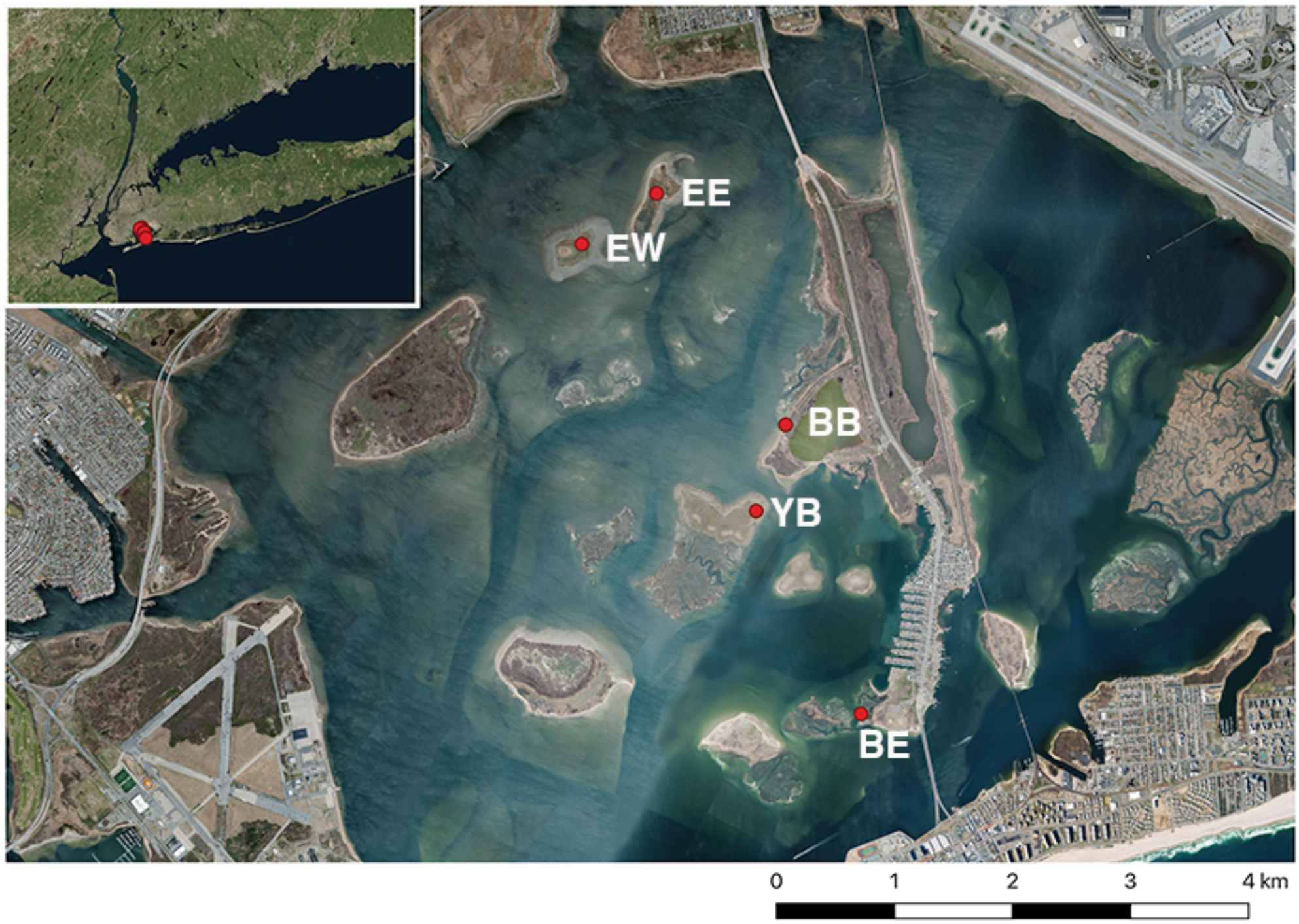
Map of sampling locations in Jamaica Bay, NY. Yellow Bar (YB), Elders West (EW), Elders East (EE), Big Egg (BE), Black Bank (BB).

## RESULTS

### Vegetation and Sediment characteristics

We found significant differences in belowground biomass among sites, but detected no clear patterns in the other vegetation variables with respect to marsh age (Table 1). Belowground biomass was greater in the older restored sites and the natural marsh site relative to the two youngest restored marsh sites (YB and EW; Table 1). Stem heights and density, leaf area, and aboveground biomass were similar among sites.

**Table 1.** Environmental variables (mean ± standard error) measured at four restored salt marshes and a natural degraded salt marsh (Black Bank) in Jamaica Bay (NY) during July 2015. Letters indicate significant differences between study sites.

| Site | n | Yellow Bar | Elders West | Elders East | Big Egg | Black Bank |
| --- | --- | --- | --- | --- | --- | --- |
| Vegetation Variables |  |  |  |  |  |  |
| Leaf Width (cm) | 10 | 0.94 (0.03) | 0.96 (0.05) | 1.06 (0.04) | 0.94 (0.06) | 0.95 (0.06) |
| Specific Leaf Area (cm<sup>2</sup> g<sup>-1</sup>) | 10 | 104.30<sup>a</sup> (2.65) | 105.95<sup>a</sup> (4.26) | 102.20<sup>a</sup> (4.98) | 134.34<sup>b</sup> (7.24) | 118.87<sup>a,b</sup> (6.26) |
| Leaf Carbon (%) | 10 | 41.78<sup>a</sup> (0.18) | 43.67<sup>b</sup> (0.35) | 42.49<sup>a</sup> (0.27) | 43.03<sup>a,b</sup> (0.31) | 42.14<sup>a</sup> (0.27) |
| Leaf Nitrogen (%) | 10 | 1.70 (0.10) | 1.87 (0.11) | 1.64 (0.07) | 1.83 (0.14) | 1.54 (0.10) |
| Avg. Stem Height (cm) | 5 | 52.0 (5.3) | 70.4 (6.3) | 75.5 (9.9) | 51.1 (7.6) | 66.6 (4.7) |
| Max. Stem Height (cm) | 5 | 91.2<sup>a,b</sup> (6.0) | 118.1<sup>b</sup> (10.0) | 109.3<sup>a,b</sup> (10.6) | 83.7<sup>a</sup> (4.8) | 91.9<sup>a,b</sup> (4.9) |
| Stem Density (m<sup>-2</sup>) | 5 | 153 (22) | 155 (30) | 211 (110) | 205 (60) | 266 (39) |
| Aboveground Biomass (g m<sup>-2</sup>) | 5 | 147 (28) | 365 (66) | 614 (265) | 202 (81) | 664 (115) |
| Belowground Biomass (g m<sup>-2</sup>) | 5 | 43<sup>a</sup> (17) | 233<sup>a</sup> (156) | 777<sup>ab</sup> (362) | 2150<sup>b</sup> (616) | 781<sup>ab</sup> (273) |
| Sediment Variables (0-5 cm) |  |  |  |  |  |  |
| Chlorophyll a (µg cm<sup>-3</sup>) | 6 | 5.78 (1.21) | 6.69 (1.63) | 8.00 (1.05) | 4.72 (0.79) | 5.67 (0.50) |
| Organic Content (%) | 6 | 0.38<sup>a</sup> (0.06) | 0.55<sup>a</sup> (0.41) | 1.13<sup>a</sup> (0.25) | 0.69<sup>a</sup> (0.11) | 15.98<sup>b</sup> (2.84) |
| Organic Carbon (%) | 6 | 0.14<sup>a</sup> (0.01) | 0.23<sup>a</sup> (0.02) | 0.53<sup>a</sup> (0.09) | 0.27<sup>a</sup> (0.04) | 7.66<sup>b</sup> (1.04) |
| Total Nitrogen (%) | 6 | 0.02 (0.01) | 0.03 (0.00) | 0.05 (0.01) | 0.06 (0.01) | 0.50 (0.08) |
| Ext. NH<sub>4</sub><sup>+</sup> (mg N L<sup>-1</sup>) | 6 | 3.82<sup>a</sup> (0.54) | 7.72<sup>a,b</sup> (0.51) | 9.32<sup>b</sup> (1.64) | 4.69<sup>a,b</sup> (0.75) | 9.49<sup>b</sup> (1.83) |
| Ext. NO<sub>2</sub><sup>-</sup> (mg N L<sup>-1</sup>) | 6 | 0.07<sup>a,b</sup> (0.01) | 0.12<sup>a</sup> (0.02) | 0.09<sup>ab</sup> (0.01) | 0.05<sup>b</sup> (0.00) | 0.07<sup>a,b</sup> (0.01) |
| Ext. NO<sub>3</sub><sup>-</sup> (mg N L<sup>-1</sup>) | 6 | 0.05 (0.00) | 0.08 (0.03) | 0.13 (0.08) | 0.03 (0.00) | 0.03 (0.02) |
| Ext. SRP<sup>-</sup> (mg P L<sup>-1</sup>) | 6 | 0.06 (0.01) | 0.12 (0.02) | 0.16 (0.07) | 0.10 (0.05) | 0.28 (0.12) |
| Sediment Variables (5-10 cm) |  |  |  |  |  |  |
| Organic Content (%) | 6 | 0.85<sup>a</sup> (0.67) | 0.37<sup>a</sup> (0.10) | 0.70<sup>a</sup> (0.10) | 1.52<sup>a</sup> (0.88) | 9.62<sup>b</sup> (1.67) |
| Organic Carbon (%) | 6 | 0.08<sup>a</sup> (0.02) | 0.15<sup>a</sup> (0.04) | 0.23<sup>a</sup> (0.03) | 0.17<sup>a</sup> (0.04) | 5.45<sup>b</sup> (1.04) |
| Total Nitrogen (%) | 6 | 0.00<sup>a</sup> (0.00) | 0.01<sup>a</sup> (0.00) | 0.02<sup>a</sup> (0.01) | 0.06<sup>a</sup> (0.01) | 0.35<sup>b</sup> (0.06) |
| Ext. NH<sub>4</sub><sup>+</sup> (mg N L<sup>-1</sup>) | 6 | 1.6 (1.0) | 2.36 (1.28) | 1.82 (0.59) | 3.02 (0.77) | 2.43 (0.21) |
| Ext. NO<sub>2</sub><sup>-</sup> (mg N L<sup>-1</sup>) | 6 | 0.09<sup>a</sup> (0.01) | 0.08<sup>a</sup> (0.00) | 0.13<sup>b</sup> (0.01) | 0.07<sup>a</sup> (0.01) | 0.07<sup>a</sup> (0.01) |
| Ext. NO<sub>3</sub><sup>-</sup> (mg N L<sup>-1</sup>) | 6 | 0.08 (0.01) | 0.04 (0.02) | 0.18 (0.09) | 0.04 (0.00) | 0.04 (0.01) |
| Ext. SRP (mg P L<sup>-1</sup>) | 6 | 0.02 (0.00) | 0.06 (0.06) | 0.02 (0.01) | 0.04 (0.01) | 0.01 (0.00) |

We observed greater sediment organic content and organic carbon content in the natural marsh (BB) in both the surface and subsurface samples, relative to restored sites. Surface sediment organic content and organic carbon were similar among the restored marsh sites, ranging from 0.38-1.13% and 0.14-0.53%, respectively (Table 1). The subsurface samples showed a similar pattern. Extractable NH_4_^+^ ranged from 3.82 to 9.49 mg l^-1^ in the surface sediment, with YB having the lowest and BB having the greatest concentrations. YB also had the lowest NH_4_^+^ concentration in the subsurface sediment. Other extractable nutrients were similar across sites (Table 1).

### Sequence Preparation

Raw sequence processing produced 594,531 paired end surface level sequences and 316,424 paired end subsurface level sequences. Surface samples had a mean sequence count of 23,671.2 (standard deviation ± 4419.0), while subsurface samples had a mean sequence of 21,904.9 (standard deviation ± 5364.7). After pooling sequences to assign amplicon sequence variants (ASVs), taxonomy was assigned using the Ribosomal Database Project classifier (27). ASVs that were classified to an unassigned Kingdom or determined to be of Archaeal, mitochondrial, or chloroplastic origin were discarded from the dataset prior to further analysis. A total of 3,887 surface ASVs and 2,569 subsurface ASVs were then used in Phyloseq (28) for community analysis.

### Alpha and Beta Diversity of bacterial communities at two sampling depths

Surface ASVs were classified into 29 phyla and 159 families, while subsurface samples were classified to 29 phyla but only 123 families. Surface and subsurface alpha diversity values were estimated using Breakaway (29) for sample richness, and DivNet (30) for Shannon Diversity (Fig. 2). Within each sampling depth, we detected no significant differences in alpha diversity among sites. However, there were significant differences between surface and subsurface samples at each site. Richness estimates ranged from 3,669 to 4,838 ASVs at the surface level, and 2,454 to 3,004 at the subsurface level. Shannon diversity estimates by location ranged from 6.14 to 6.86 at the surface level and 5.17-6.25 at the subsurface level. Minimum and maximum richness estimates and Shannon estimates are provided by location and sampling depth in Table S1.

**FIG 2.**
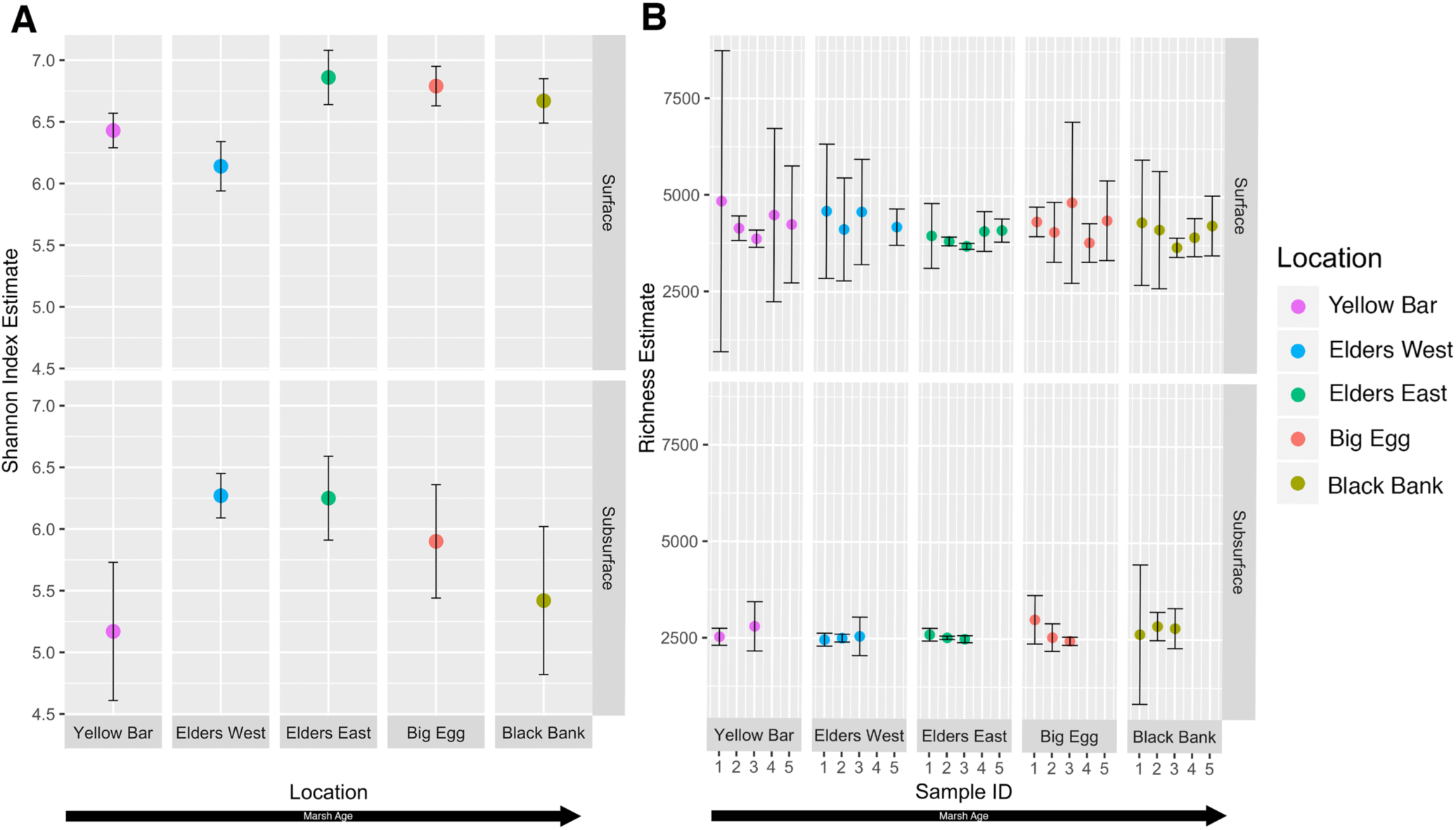
Alpha diversity estimates displayed by marsh (A and B) and sampling depth (surface and subsurface). (A) Shannon entropy estimates with confidence intervals made with DivNet, site wide estimations made at the ASV level. (B) Richness estimates with confidence intervals using Breakaway. Two outliers EWS4 and YBD2 were excluded from the richness plot.

Unweighted and normalized weighted UniFrac (31) distances were calculated using Phyloseq, and differences in community structure were assessed with NMDS ordinations in R (32) (Fig. 3). PERMANOVA analysis using Bray-Curtis distances were conducted using Vegan (33) and showed that sites were grouped significantly by location at the surface level but not at the subsurface level (Table 2).

**Table 2.** PERMANOVA results of surface and subsurface sediment microbial communities in relation to the environmental variables.

| **Surface Sediment<sup>a</sup>** | **p-value** | **R<sup>2</sup>** |
| --- | --- | --- |
| Belowground biomass | 0.597 | 0.016 |
| Organic content % | 0.361 | 0.021 |
| Organic C % | 0.517 | 0.018 |
| Total N % | 0.621 | 0.016 |
| Ext. NH<sub>4</sub><sup>+</sup> | 0.433 | 0.020 |
| Ext. NO<sub>2</sub><sup>-</sup> | 0.438 | 0.019 |
| Ext. NO<sub>3</sub><sup>-</sup> | 0.519 | 0.018 |
| Ext. SRP | 0.597 | 0.016 |
| Location | 0.002*** | 0.192 |
| **Subsurface Sediment<sup>b</sup>** | **p-value** | **R<sup>2</sup>** |
| Belowground biomass | 0.803 | 0.043 |
| Organic content % | 0.902 | 0.035 |
| Organic C % | 0.967 | 0.028 |
| Total N % | 0.983 | 0.025 |
| Ext. NH<sub>4</sub><sup>+</sup> | 0.503 | 0.067 |
| Ext. NO<sub>2</sub><sup>-</sup> | 0.852 | 0.038 |
| Ext. NO<sub>3</sub><sup>-</sup> | 0.938 | 0.032 |
| Ext. SRP | 0.822 | 0.041 |
| Location | 0.793 | 0.207 |
| <sup>a</sup> n=5 |  |  |
| <sup>b</sup> n=3 |  |  |

**FIG 3.**
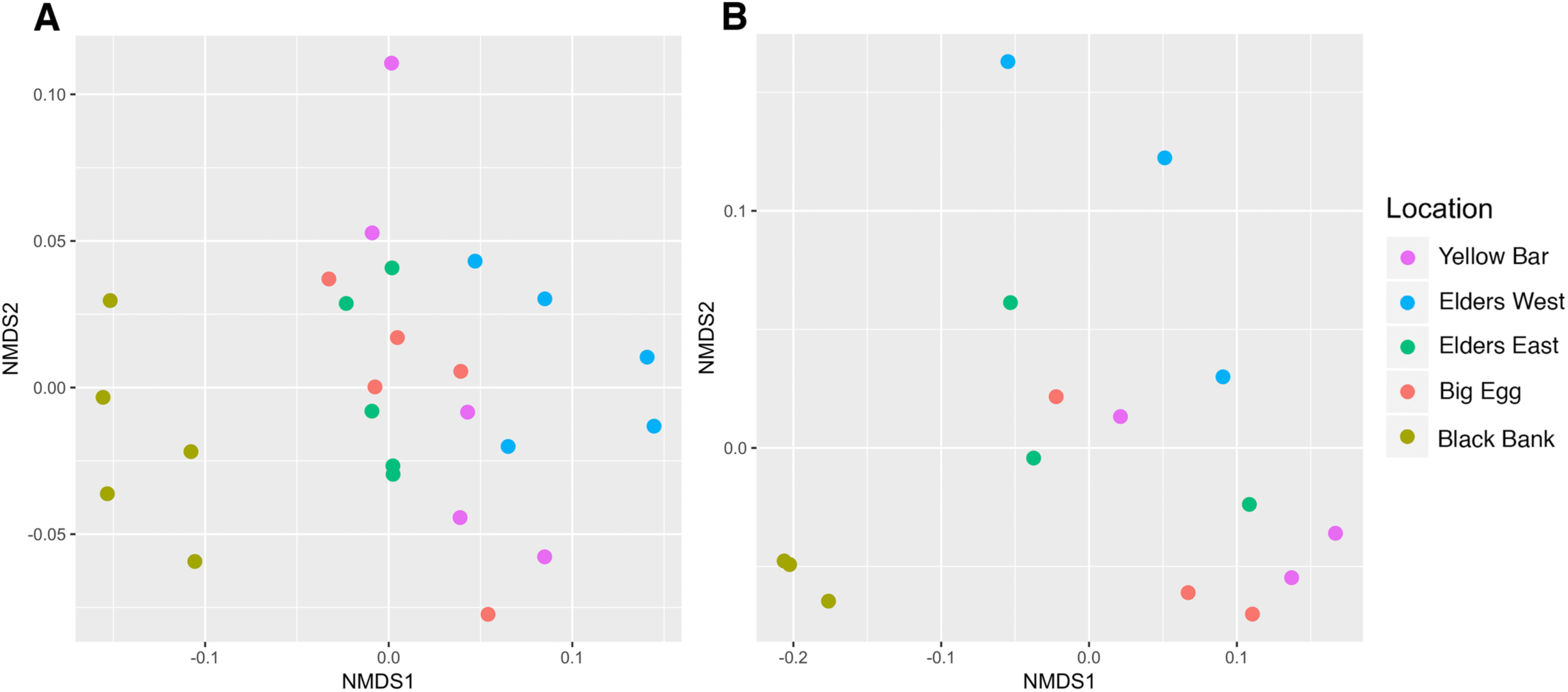
Non-metric multidimensiona scaling (NMDS) of weighted UniFrac distances. Prior to UniFrac calculations ASV counts were variance stabilized using DESeq2. Surface samples (A) significantly grouped by location in PERMANOVA analysis (p=0.001*** R^2^=0.301), while subsurface samples (B) grouped non-significantly by location (p=0.151, R^2^=0.258).

Hierarchical clustering was performed to visualize weighted UniFrac sample distances. UPGMA trees were constructed for both surface and subsurface samples (Fig. 4). At the surface level, BB clearly separates from all other locations, while younger restored EW and YB marshes are grouped together, and older restored EE and BE marshes are grouped together. Subsurface samples show another distinct separation of BB with less distinct clustering of the other sites.

**FIG 4.**
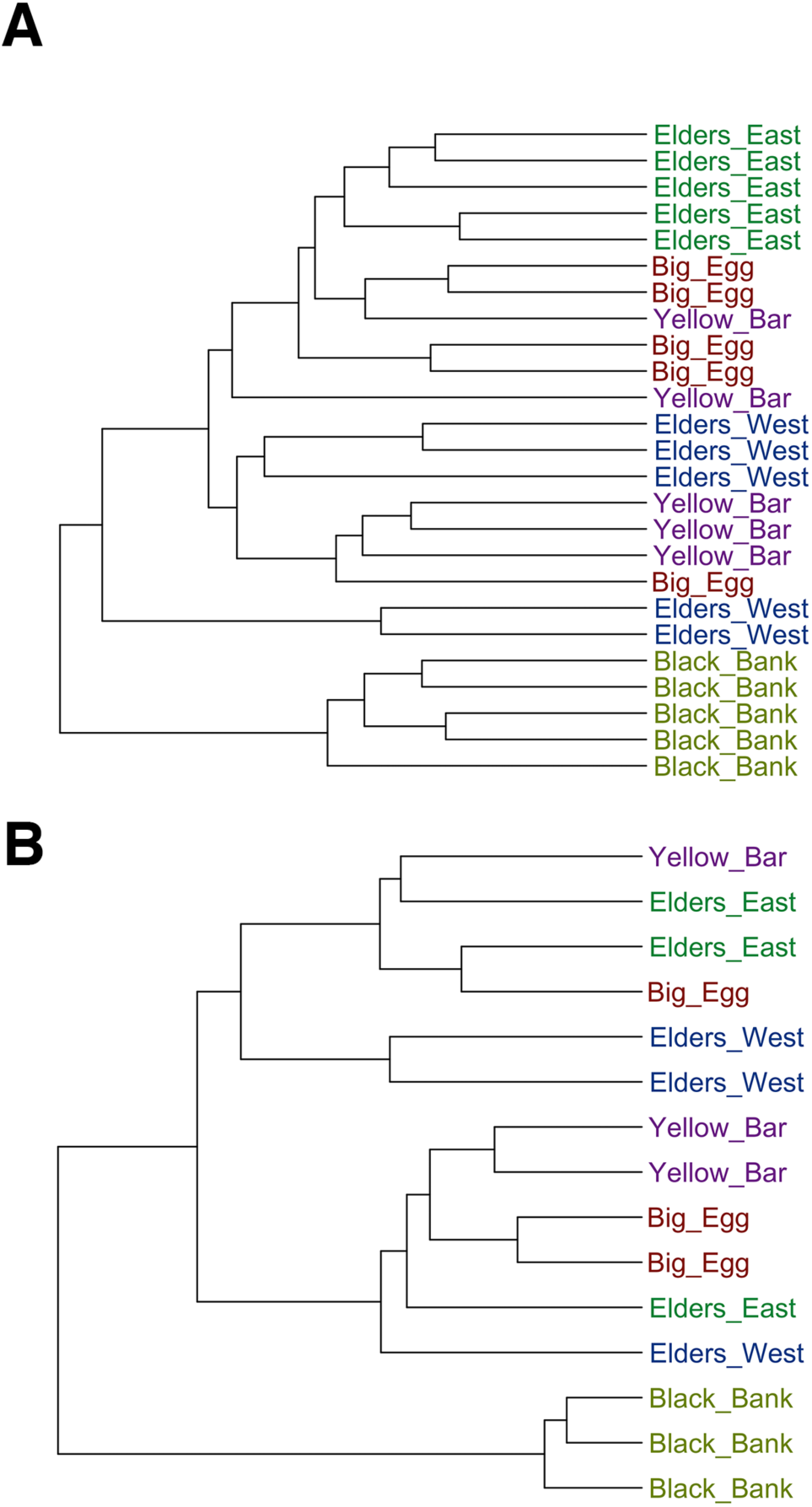
UPGMA dendrograms of weighted UniFrac distances of surface (A) and subsurface (B) sediment communities. Prior to UPGMA clustering, ASV tables were variance stabilized using DESeq2.

### Taxonomic Composition by relative abundance

Fig. 5A-B shows the taxonomic composition at the phylum level of all ASVs found at greater than 1% relative abundance. For all samples and at both sampling depths, *Proteobacteria* was the most dominant phylum of the community, with *Bacteroidetes* second. Seven different phyla were found to be differentially abundant between restored and unrestored locations at the surface level using DESeq2 (34) or displayed clear trends with age (Fig. 5C).

**Fig 5.**
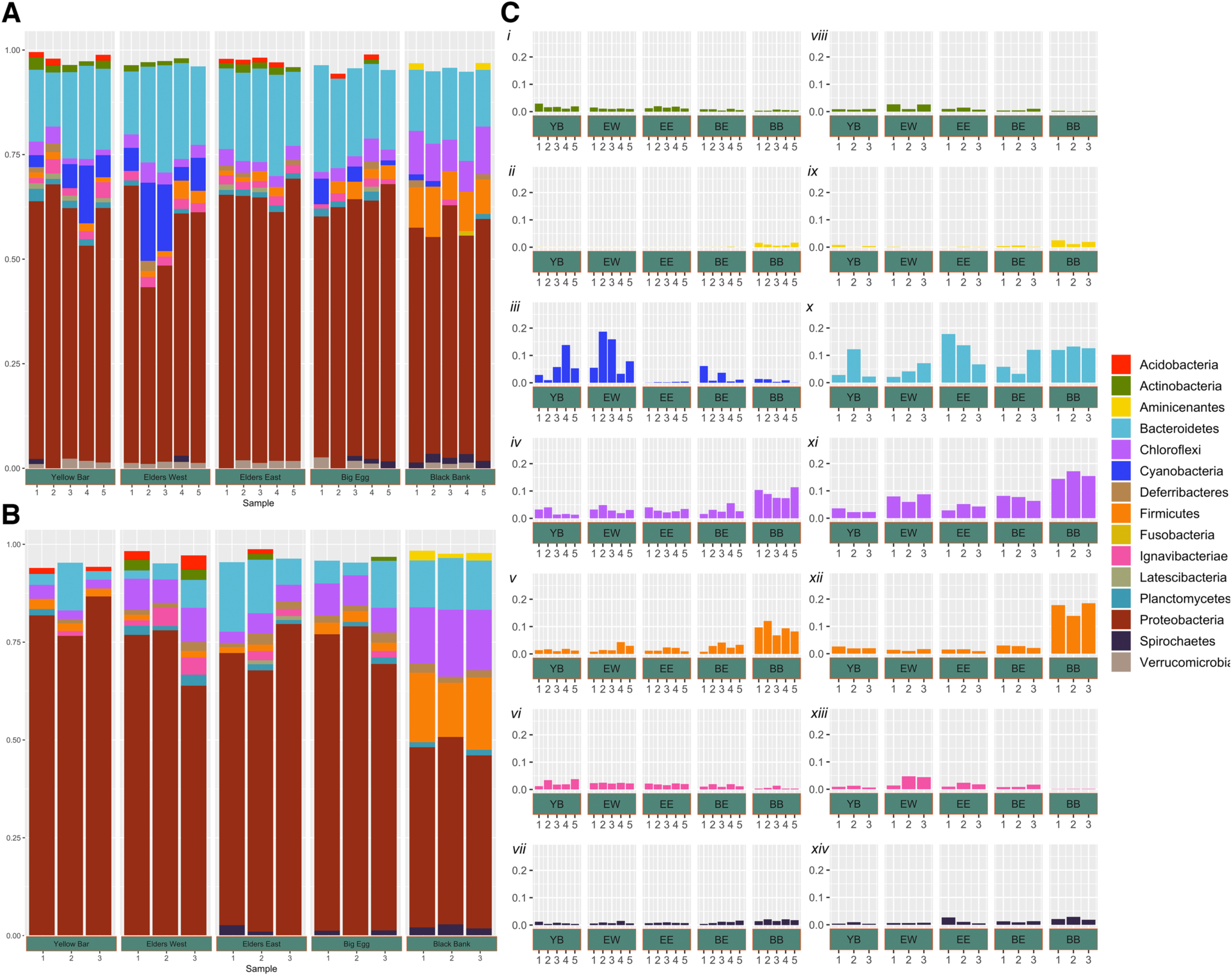
**Taxonomic composition** in relative abundance at the phylum level of sediment bacterial communities from the surface (A) and subsurface (B) and grouped by sampling marsh. Only phyla representing >=1% of the total community are displayed (A) and (B). Individual phyla from the surface (*Ci-Cvii*) and subsurface (*Cvii-Cxiv*) that display trends across the chronosequence or are differentially abundant between restored and unrestored.

### Surface Taxonomy

The relative abundance of the phylum *Chloroflexi* was found to be highest at BB (mean ± sd: 9.1 ± 1.8%) compared with all other sites (2.8 ± 1.1%). Within *Chloroflexi, Anaerolineaceae* was the dominant family at the restored sites, while BB was roughly equally split among *Anaerolineaceae, Dehalococcoidetes*, and *Chloroflexi*, which could not be classified to below the phylum level.

*Firmicutes* was also more common (9.2 ± 1.9%) at BB than any other site (2.0 ± 1.1%). At the class level the composition of the *Firmicutes* phylum differs among sites. All restored sites were dominated by *Clostridia* except for BE. At BE *Firmicutes* was roughly split between *Clostridia* and unknown *Firmictutes*, while BB was dominated by unknown *Firmicutes*.

*Cyanobacteria* were more common at the two youngest sites, EW (10.3 ± 6.7%) and YB (5.7 ± 4.9%), relative to EE, BE or BB (0.2 ± 0.1%; 2.4 ± 2.4%; 0.8 ± 0.6%, respectively). As post-restoration marsh age increased, so did the relative abundance of *Spirochaetes* (0.7 ± 0.3% to 1.8 ± 0.3%), dominated by *Spirochaeta*. The relative abundance of *Actinobacteria* and *Ignavibacteriae* decreased with marsh age (1.9 ± 0.7% to 0.5 ± 0.2% and 2.4 ± 1.1% to 0.6 ± 0.4%, respectively). Among sites the families that make up *Actinobacteria* varied, while *Ignavibacteriae* was dominated by *Ignavibacterium* at all sites.

### Subsurface Taxonomy

*Chloroflexi* and *Firmicutes* were also more common in subsurface samples at BB (15.6 ±1.4% and 16.7 ± 2.5%) than at any other site (5.4 ± 2.4% and 1.8 ± 0.7%). As with surface samples, the contributions of *Actinobacteria* and *Ignavibacteriae* to the total community decreased with marsh age EW to BB (2.1 ± 1.0% to 0.2 ± 0.06% and 3.5 ± 1.8% to 0.1 ± 0.02%). YB, however, did not follow this trend (0.8 ± 0.1% and 0.9 ± 0.3%). *Actinobacteria* were dominated by the class *Actinobacteria*, and *Ignavibacteriae* were solely *Ignavibacterium*. The phylum *Aminicenantes* contributed more to the community at BB (1.8 ± 0.7%) than any other site (0.2 ± 0.3%), while *Acidobacteria* were more prevalent at the two youngest sites with 2.3 ± 1.3% at EW and 1.1 ± 0.3% at YB compared to BB (0.4 ± 0.07%), BE (0.7 ± 0.2%) and EE (0.8 ± 0.4%). Finally, the phylum *Bacteroidetes* made a larger overall contribution to community structure at the two oldest restored sites EE (12.7 ± 5.6%) and BE (7.0 ± 4.5%), and BB (12.6 ± 0.6%) compared to the two youngest restored sites EW (4.4 ± 2.5%) and YB (5.7 ± 5.6%).

### Constrained Analysis of Principal Coordinates (CAP)

We used constrained ordinations to explore how environmental variables (Table 1) were associated with changes in community composition at both the surface and subsurface depths. At the surface level, the main environmental factors associated with the separation of BB from all other sites were belowground biomass, and the percent of total N, organic C, and organic content found in sediments (Fig. 6). The separation of Elders West from other restored sites was associated with differences in extractable NO_2_^-^ and NO_3_^-^. At the subsurface level, percent total N, organic C, and organic content correlated with the separation of BB from all other samples, whereas belowground biomass did not. Subsurface EE and EW were separated from BE and YB, and this difference in community composition was associated with extractable NO_2_^-^, NH_4_^+^, soluble reactive phosphate (PO_4_^3-^) and sediment organic content.

**FIG 6.**
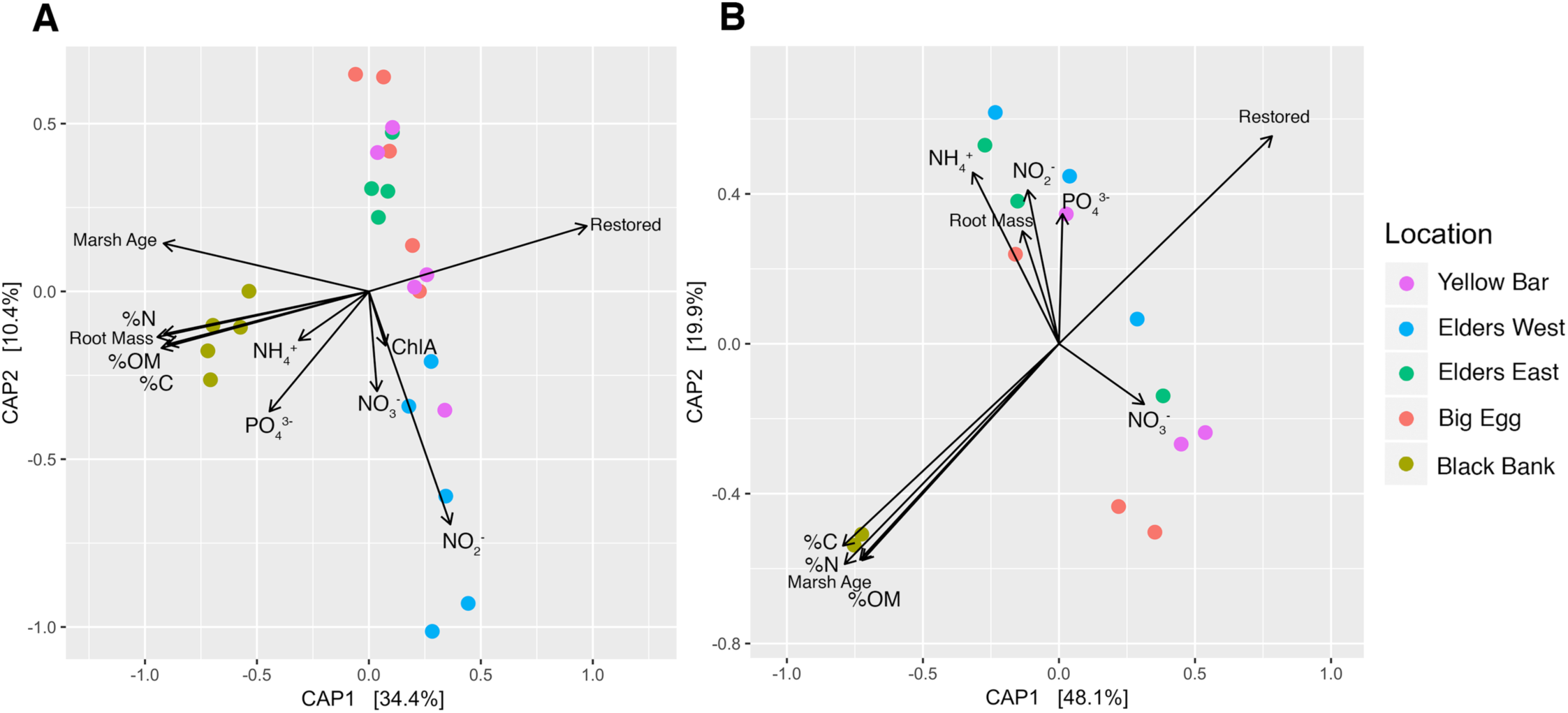
**Canonical analysis of principal coordinates (CAP)** ordinations of weighted UniFrac distances. Bacterial communities of surface (A) and subsurface (B) sediment are colored by sampling location. Fitted vectors of environmental variables represent the direction of the gradient and the length represents the strength of the variable.

In addition, regressions were made on NMDS and CAP axes weighted by the inverse of the standard error of the predictor variable (Fig. 7), to assess their effect on bacterial community composition. Black Bank separates out based on sediment organic content, but the trend in community structure in the restored marshes appeared more related to belowground biomass.

**FIG 7.**
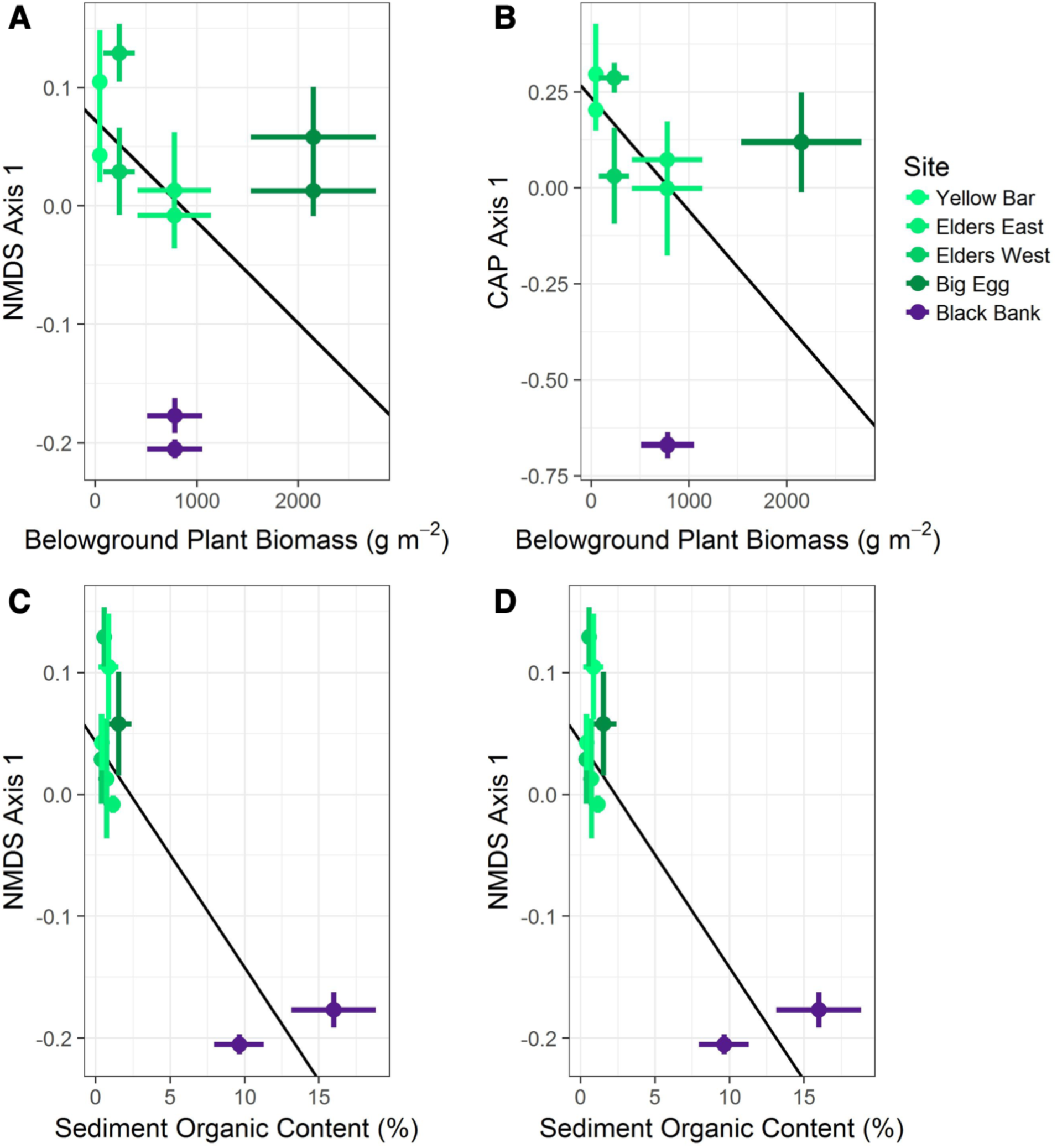
**Weighted regressions** using environmental factors as predictor variables weighted by SE.

### Predicted nitrogen cycling gene content

ASV sequences were run through the PAPRICA (35) metabolic inference software to predict the abundance of genes involved in N cycling. After rarefaction to account for uneven sequencing depth and correcting for 16S gene copy number, bar graphs of predicted gene abundances were constructed to show differences among sampling sites and sampling depths (Fig. 8). Spearman correlation coefficients were calculated to determine which environmental variables exhibited positive and negative correlations with predicted N gene content (Table 3).

**Table 3.**
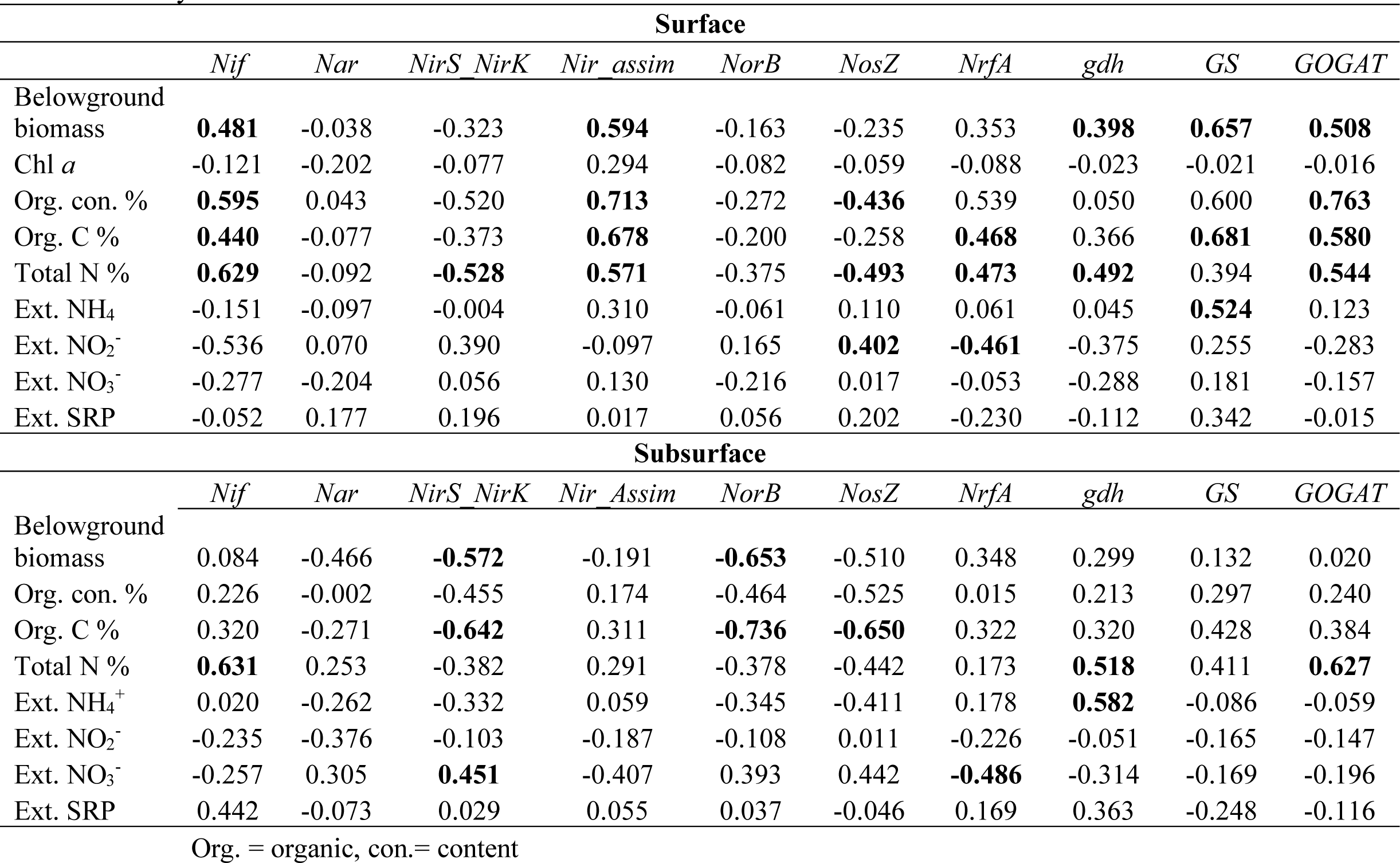
Spearman correlation coefficients of the relationships between environmental variables and the predicted nitrogen cycling gene abundances from the surface (0-5cm) and subsurface (5-10cm) sediment samples. Statistically significant relationships are indicated by bold font.

**FIG 8.**
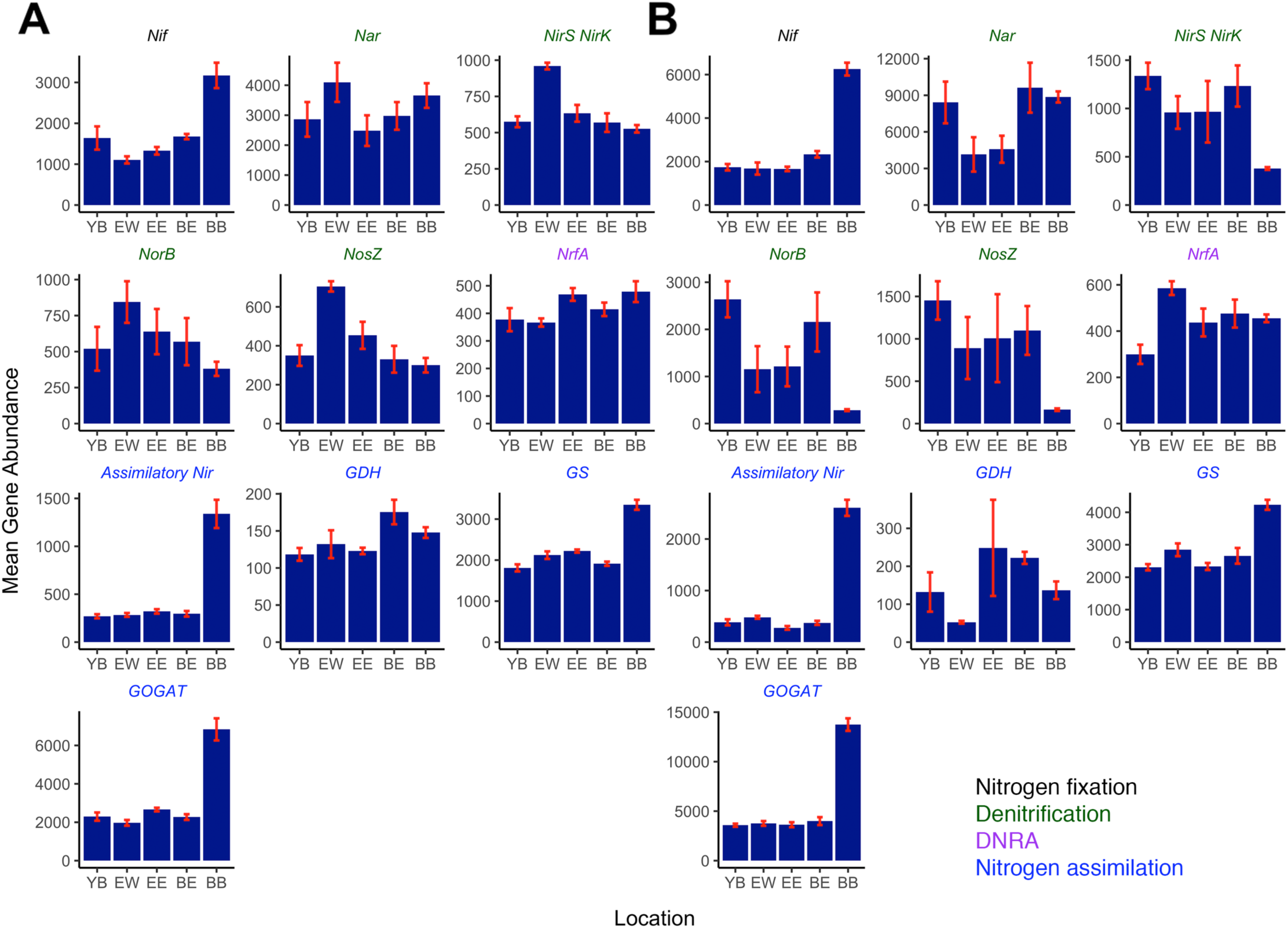
Predicted nitrogen cycle gene abundances by location (Yellow Bar=YB, Elders West=EW, Elders East=EE, Big Egg=BE, Black Bank=BB) and sediment depth. Nitrogen fixation (Nif), denitrification (NirS, NirK, NorB, NosZ), DNRA (NrfA), and nitrogen assimilation (assim. Nir, GDH, GS-GOGAT) were predicted for both surface (A) and subsurface (B) sediment communities using PAPRICA. Prior to gene prediction sequences were randomly subsampled to even depth across all samples at both sampling depths.

At the surface level, predicted nitrogen fixation gene (*Nif*) counts increased with marsh age and were most abundant at BB. Genes involved in denitrification (*Nar, NirS, NirK, NorB, NosZ*) were highest at the EW site. We did not find a clear trend in genes associated with DNRA (*NrfA*) among sites. Assimilatory nitrite reductase (*Nas*) was significantly more abundant at BB but similar across all other sites, as were genes involved with ammonia assimilation (*gdh* and the GS-GOGAT pathway).

Subsurface samples from YB and BE showed the most predicted denitrification genes, and the abundance of these genes was roughly double compared with surface samples. Similar patterns were predicted in subsurface genes involved in assimilatory nitrite reduction, DNRA and GS-GOGAT as in the surface. *Nar* was more abundant at YB, BE, and BB, and *NorB* was highest at YB and BE.

## DISCUSSION

### Complex system, complex communities

Salt marsh sediments are complex systems with diverse microbial communities that perform a wide array of biogeochemical processes critical to the world’s nutrient cycles and the global transfer of materials between terrestrial and ocean ecosystems. While much is known about the processes that microbial communities are able to perform as a whole in these systems, more research is needed to elucidate the specific taxonomic and functional composition of these communities, and the environmental drivers that shape them. Here our results showed that five marsh islands within the same estuary exhibited distinct differences in Shannon diversity, as well as beta diversity, taxonomic composition, and predicted functional capacity among sampling sites. These differences were also observed when comparisons were made between surface and subsurface samples within the same sampling location, attempting to tease out differences in community composition under different redox conditions. Some differences in community structure among sites and depths can be correlated to the environmental data, or known functional characteristics of taxa. Other results, however, do not have a clear explanation, and may be influenced by daily inundation times, aspects of restoration such as sediment origin, or variables not measured such as hydrology or erosion.

### Alpha diversity estimates differ by both location and sampling depth

Richness estimates at all five sampling sites showed higher richness at the surface than the subsurface level. Within these sampling depths, however, we found no significant differences in richness across the chronosequence. Similarly, a salt marsh chronosequence spanning over 100 years showed no age-related differences in richness (15). The observed differences between sampling depths in the current study could result from both more stringent selective pressures at the subsurface level than at the surface level, as well as less frequent perturbation and taxonomic turnover at subsurface levels. Sediment oxygen conditions are typically a major driver of community composition and could restrict the number of taxa able to persist at the subsurface level.

Estimates of Shannon diversity, which take into account evenness among taxa, showed significant differences among the five marsh sites at both surface and subsurface levels, consistent with our hypothesis. While we found no strong trends in diversity with age or restoration status across sites, the two youngest sites had the lowest Shannon diversity. We hypothesize that these younger restored marshes may be farther away from ecological equilibrium due to their sediment composition or nutrient limitations.

Alpha diversity estimation in microbiome studies is an area of active research, and values may be under- or overestimated due to the compositional nature of the data (36, 37). However, as long as no biases exist across samples, Shannon diversity remains useful for exploration of broad relative trends. In this study we note that environmental factors may be driving differences in alpha diversity among sites and between depths. In a separate study shannon diversity and richness were found to differ between sampling depths in mudflat sediment (38). However, trends described in this study were not clear as two of three sites showed higher richness at subsurface depths, and two of three sites showed lower Shannon diversity at the subsurface (38). A separate study calculated richness and Shannon estimates based on nitrogen functional gene diversity, and found that diversity decreased with marsh age across a 100 year time span (16), much longer than the 12 year span in this study.

### Community composition differs significantly among marsh sites

Beta diversity calculations from surface sediment demonstrate significant groupings by location, despite being proximate geographically (<4.6 km between the two most distant sites). This result was consistent with the findings of Dini-Andreote et al. (15) which looked at a 100 year naturally occuring chronosequence. In contrast, significant groupings were not observed at the subsurface level. The unrestored and degrading site (BB) was distinct from the other locations, with percent N, C, organic content, and belowground plant biomass positively correlated with this separation. Differences in organic matter accumulation and belowground biomass have been observed between degrading and restored marshes previously in Jamaica Bay and elsewhere (21, 39). These differences suggest that unrestored marshes harbor distinct microbiomes when compared to those from newly constructed marshes that may be related to belowground root growth, decomposition, or greater accumulation of recalcitrant forms of carbon in older marshes. Newly constructed marshes are likely to differ from natural marshes in the amount of accumulation of belowground plant biomass and organic carbon. As a consequence, new plant growth and labile carbon may provide more important sources of energy for microbes in newly constructed marshes. However, an important caveat is that we are unable to determine with our dataset how this difference may be affected by the degraded status of Black Bank. We expect degraded marshes to show less new root growth and a higher fraction of standing root mass that consists of recalcitrant (*i.e.*, difficult to decompose) material, relative to healthy ones. Future studies should incorporate comparisons of stable and degrading natural marshes wherever possible in order to determine how marsh degradation affects these factors. While the factors separating BB from the restored marshes were relatively clear, surprisingly, neither belowground biomass nor percent organic content, N, or C, had a statistically significant effect on groupings from PERMANOVA analysis. Therefore factors driving statistically distinct communities among the restored marsh islands may not have been captured in our environmental data.

An UPGMA dendrogram of surface level communities (Fig. 4A) shows that microbial communities of EE and BE are more similar despite being the sites most geographically distant from each other (Fig. 1). EW and YB were also similar to each other, and BB was the most dissimilar to the other four sites. While we cannot draw firm conclusions from this clustering due to the complex nature of the study system and limited sample size, the clusters share some distinctive characteristics. EE and BE are the two oldest restored marshes, and both have demonstrated loss of plant cover post-restoration between 2008 to 2012 (23). EW and YB are the two youngest restored marshes, and likely retain a higher elevation relative to the older restored sites. They have also gained more plant cover from 2008 to 2013 (24). Consistent with our predictions, the microbial community at the unrestored site, BB, was clearly dissimilar from all other sites. These differences could be related to marsh elevation as other studies (21, 40) have demonstrated that increased inundation may alter sediment and vegetation characteristics (21, 40). Changes in patterns of inundation, vegetation, and sediment traits would likely affect microbial communities as well.

In the subsurface UPGMA dendrogram (Fig. 4B), BB communities were also dissimilar in composition, but communities from the other four sites did not cluster by location. Restored marshes have higher percentage of sand and accumulate less organic matter and total N in subsurface layers (21, 39). Nutrient availability and sand content have previously been identified as significant drivers of salt marsh microbial community structure across a chronosequence (15, 16). It is possible that subsurface sediments experience less site-to-site variability in environmental conditions than surface sediments, leading to slower divergence in community composition at subsurface depths. As a consequence, bacterial communities below the surface may change more slowly than at the surface following restoration. Several studies have shown that microbial community structure, functional potential, and nitrogen cycling processes change across a natural marsh chronosequence (15–17, 41); however to our knowledge post-restoration succession within the microbial community has not been examined previously at the subsurface level. More research is needed to determine if these results are due to a difference in the rate of community succession between surface and subsurface, or if subsurface microbial communities are similar among unrestored marsh sediment as well.

### Taxonomic composition shows distinct differences among sampling sites

The two most dominant phyla, *Proteobacteria* and *Bacteroidetes*, observed at all study sites were also most dominant in other studies performed in estuarine systems (15, 42–44). Several other phyla showed striking variation at both surface and subsurface samples across sites, which was consistent with our hypothesis that microbial communities would differ among sites along the chronosequence. The phylum *Chloroflexi* was relatively more abundant at BB than any other site at the surface and subsurface levels. The dominant family within *Chloroflexi* was *Anaerolineaceae* in restored surface sites, similar to findings from a previous study in mudflat sediment (38). This family specifically, as well as *Chloroflexi* as a whole, have been shown to contain a wide array of carbohydrate hydrolytic genes (45), which correlates with the high amount of accumulated organic matter and decomposing belowground biomass at BB (21), and aligns with CAP results showing organic content and C to be positively associated with the BB site. This observation accords with our third hypothesis that communities in older marshes would harbor more taxa with the metabolic capacity to break down organic matter. Previous studies have also shown that *Chloroflexi* was significantly more abundant in older marshes across a natural chronosequence (15), and was associated with N- and organic-rich sediments (15, 43).

*Firmicutes* was another phylum more abundant at both the surface and subsurface level at BB. In contrast, a previous study showed *Firmicutes* to be associated with younger sites in a natural chronosequence (15). Unlike *Chloroflexi, Firmicutes* was dominated by unclassified *Firmicutes* in our results, and the proportions of classified families within the *Firmicutes* phylum vary from site to site. *Firmicutes* could represent a phylum that responds to site-specific differences in environmental parameters, where certain families can outcompete others. Alternatively, metabolic capacities within *Firmicutes* may be similar enough that different families may fill the same metabolic role at different sites. More research is needed to determine what role each *Firmicutes* ASV is filling in this system.

At the surface level, *Cyanobacteria* were major contributors to overall composition at YB and EW, the two youngest sites. These cyanobacteria were likely filling the role of autotrophic N fixers, consistent with our second hypothesis. This result is consistent with studies previously conducted in marsh chronosequences (15, 17) and is likely due to the ability of many *Cyanobacteria* to fix atmospheric nitrogen as early colonizers of marsh communities and that the youngest sites are likely to be nitrogen-limited (46, 47), which is supported by our nutrient analysis results (Table 1).

At the subsurface level, *Bacteroidetes* were relatively more abundant at BB, BE, and EE (unrestored and two oldest) when compared to EW and YB (two youngest). This difference could be due to accumulation of organic matter over time and the previously shown characteristic of *Bacteroidetes* phylum to have a broad array of carbohydrate-degrading genes (45). A recent study, however, examined diversity at the surface level of a natural marsh chronosequence, and found *Bacteroidetes* to be associated with younger sites (0 and 5 years of age) (15), possibly indicating that *Bacteroidetes* fulfills a different role in subsurface samples than at the surface.

### Predicted nitrogen cycling gene content

Nitrogen fixation predicted gene content (nitrogenase, *Nif*) increased slightly across the chronosequence. Gene content in surface sediment at the unrestored site (BB) was substantially higher, with twice as many predicted *Nif* gene copies than any other site, and three times higher counts in subsurface samples. This result was surprising as we predicted *Nif* gene copies would be greatest at the young restored marshes. Interestingly, we did observe a higher abundance of *Cyanobacterial* autotrophic N fixers at EW and YB. One possible explanation for this is that *Cyanobacteria* are underrepresented in the pre-constructed reference genome database used to calculate our gene predictions (9 genomes out of ∼5400). However, this increase in *Nif* gene content with age, despite there being a lack of fixed N at younger sites, has been shown in one previous study (16) through qPCR analysis of the *Nif* gene, a method independent of genome databases. The chronosequence of marshes in that study spanned ∼100 years, somewhat shorter compared with the estimated age of Black Bank (>200 years). Alternatively, the higher gene content at BB and the older restored marshes could be due to heterotrophic N-fixing anaerobic bacteria that use sulfur for respiration (48), rather than cyanobacteria. This possibility is consistent with our observation that surface samples had less predicted *Nif* content than the subsurface. Significant positive relationships between predicted *Nif* content and percent N at the surface, and with belowground biomass, percent organic content, C and N in subsurface samples (Table 3), suggest an increase in heterotrophic N fixation at mature marshes where C content is higher. This would also be consistent with other studies that report high sulfide levels in sediment with greater accumulation of recalcitrant carbon and marshes experiencing greater inundation (*i.e.*, lower marsh elevation) (21, 48).

Our results differ from a study of a naturaly developed marsh chronosequence, consisting of two young marshes (7 and 15 years) and one mature marsh (150 years) in Virginia. In that study, N fixation rates were found to be negatively correlated with organic content and were higher at younger marshes (17). An increase in *Nif* gene content with a hypothesized decrease in N fixation rates could be explained by a disconnect between the *Nif* gene presence in the predicted metagenome of BB and the actual rate of N fixation. The method presented here reports predicted gene content, rather than a direct reflection of genes being actively expressed. Many of the genomes containing *Nif* genes may be facultative N fixers who rarely fix N. Preliminary evidence has shown that denitrification was the dominant N_2_ flux at BB and that these gene copy results were likely due to a mismatch between predicted *Nif* content and actual rates of N fixation (49).

We expected to see an increase in denitrification gene predictions at the natural unrestored marsh (BB) since a previous study showed significant increases in denitrification gene content in a marsh creek that had experienced extensive nutrient loading (50). Jamaica Bay has been experiencing increased N loading for >110 years (27). Contrary to our expectation, the gene predictions demonstrated that BB microbes had less denitrification gene content than any restored marshes. Gene predictions related to the process of denitrification (*Nar, NirS/NirK, NorB, NosZ*) were lowest for most genes at BB, in both surface and subsurface samples. Surface gene content for these genes was highest at a relatively young marsh (EW). Subsurface sample counts for these genes were high overall, with the exception of BB, and were roughly two-three times higher than surface sample counts, likely due to the anaerobic conditions in the subsurface sediment. The subsurface *NirS/NirK, NorB*, and *NosZ* were significantly negatively correlated with belowground biomass and percent C. BB did display a higher abundance of genes related to nitrogen assimilation (assimilatory *Nir*, gdh and *GS-GOGAT*) than any other site at both surface and subsurface, consistent with previous work that showed an increase in nitrogen assimilation gene content with marsh age (41). This could indicate that nitrogen assimilation is favored over nitrogen removal at BB. Anaerobic sediment and high sulfides may inhibit coupled nitrification-denitrification thus reducing denitrification potential at BB. Again, these results reflect predicted gene content rather than gene expression, and it is important to confirm these findings using techniques such RNA-Seq or proteomics to give a more accurate representation of the metabolic processes occurring in these communities. Preliminary results from these locations indicate significantly higher rates of microbial denitrification at BB relative to restored sites (51). As with our nitrogen fixation results, we expect that many denitrifying bacteria are facultative. In that case, local environmental conditions that favor expression of denitrification genes may prove to be better predictors of actual N-removal rates than characteristics of the microbial community itself.

Predicted *NrfA* counts, which correspond to DNRA, showed significant positive correlations with percent organic content, C and N and a significant negative correlation with NO_2_^-^ in the surface sediment. No significant correlations with *NrfA* were found in subsurface samples (Table 3). Although we did note that younger marshes had slightly less *NrfA*, no strong trend was observed in predicted DNRA potential (*NrfA*) across the chronosequence at either sampling depth. Previous work based on full metagenomes demonstrated a decrease in relative abundance of *NrfA* genes across a chronosequence (41), unlike the predicted slight increase found here. This discrepancy may result from possible shortcomings of functional gene prediction based on 16S rRNA gene abundance compared to shotgun metagenomics, as bacterial genomes from environments such as salt marsh sediment remain relatively underrepresented in reference databases. It is also possible that our chronosequence does not span enough time to display the same trend.

## CONCLUSIONS

Our results indicated distinct differences in bacterial communities, and their metabolic capacities, among five marsh islands despite their close geographic proximity within the same estuary. The two youngest sites showed an increased relative abundance of taxa known to be early colonizers of salt marsh sediment and known autotrophic N fixers. Restored sites contained more predicted genes related to the ability to perform denitrification. The most dissimilar community at both surface and subsurface sediment depths was Black Bank, the oldest and the only unrestored marsh. Analysis of predicted gene content suggested that metabolic capacity for N assimilation was greater at BB than the restored marsh sites. We speculate that this unique community composition at BB was driven by environmental factors at this site including high C, N, and organic content. We cannot determine with the present dataset, however, whether marsh age, its degrading nature, or its unrestored status are the proximate causes for these differences. Many N processes are facultative, especially denitrification, so measuring gene expression should be considered as a replacement for the measurement of gene abundance. Future work within this and similar systems should seek to experimentally link the biogeochemical processes ongoing within the marsh to specific bacterial taxa through RNA-seq, proteomics, metabolomics and should also include stable natural marshes in addition to degrading natural marshes.

## MATERIALS AND METHODS

### Study site sediment and vegetation characteristics

Data was collected from 5 *Spartina alterniflora* salt marshes located in the center of Jamaica Bay (New York City, NY, USA). Four of these marshes were restored marshes while the fifth was a natural degrading marsh (22). Yellow bar (YB restored 2012), Elders East (EE restored 2006), Elders West (EW restored 2010), Big Egg (BE restored 2003), Black Bank (BB natural degrading; Fig. 1). Sampling was performed on 7-8 July 2015. At each site a transect was established parallel to the marsh edge (∼1m from marsh edge). A 0.25m^2^ quadrat was haphazardly tossed along the transect to collect five replicate samples from each site. Samples from each quadrat were taken for aboveground biomass by harvesting all plant material inside the quadrat. Measurements of stem density, heights, and leaf characteristics were performed in the laboratory. The belowground biomass was sampled by taking a core (2.76 × 15cm) from the center of the quadrat to a depth of ∼10cm. The entire core was brought back to the lab where it was wet-sieved through a 1mm mesh. Both above and belowground plant material was dried at 60°C and sub-samples were homogenized with a mortar and pestle. The carbon and nitrogen content of the plant material was measured using a Perkin Elmer Series 2400 CHN element analyzer (Perkin Elmer Inc., Shelton, CT) using acetanilide as a standard.

Sediment samples were collected from the belowground biomass core. Subsamples from the sediment surface (0-2.5cm) were taken to measure sediment chlorophyll-a using the acetone extraction method and measured spectrophotometrically (52). Subsamples were also taken from both the surface 0-5 cm and subsurface (5-10cm) to determine the organic content, organic carbon, total nitrogen, and extractable nutrient concentrations. Sediment was dried at 60°C and weighed to determine water content. The sediment organic content was determined based on the loss on ignition at 500°C (53). Dried sediment samples were treated with 25% hydrochloric acid and re-dried (54) to determine percent organic carbon and total nitrogen. Lastly, we used a fresh ∼5g sediment sample to determine extractable nutrient concentrations. Samples were extracted with 10-mL 2N KCl and analyzed for ammonium, nitrite, nitrate, and soluble reactive phosphate concentrations using a Seal AQ2^+^ discrete autoanalyzer (Seal Analytical Inc., Mequon, WI) following the methods of (55–57), respectively. A one-way ANOVA was used to compare environmental variables across sites. A Tukey post-hoc analysis was used to determine differences among sites. Analyses were performed with SigmaPlot 11 (Systat Software Inc., UK).

### DNA Collection and Extraction

Sediment was collected from the sediment core samples described above. Sediment was collected using a sterile scoopula (Fisher Scientific) into a sterile 50 mL Falcon tube from the surface (0-1cm) and subsurface (5-10 cm). Sediment samples were placed on ice and transported back to the lab and were frozen (−20°C) until extraction. Sediment was thawed on ice upon removal from freezer then transferred to a weigh boat using a sterile scoopula. Sediment was homogenized and approximately 0.4 g of wet sediment was used in the PowerSoil DNA isolation kit (MoBio USA) following manufacturers protocol. Extracted sediment DNA was quantified using a Nanodrop 2000 system to check for DNA concentration and purity.

### 16S Ribosomal RNA Gene Amplification

Five surface samples and three subsurface samples from each site were chosen for amplification based on DNA concentration and purity as determined by Nanodrop 2000. Amplification of the V4 variable region of the 16S small ribosomal RNA gene was completed using 515F/806R (58) primer set with a barcode on the forward primer, using a 30 cycle PCR and HotStarTaq Plus Master Mix Kit (Qiagen, USA) with the following conditions: 94°C for 3 minutes, followed by 28 cycles of 94°C for 30 seconds, 53°C for 40 seconds and 72°C for 1 minute. A final elongation step at 72°C for 5 minutes was performed. Amplified DNA sequences were pooled in equal proportions, and then purified using Ampure XP Beads, purified PCR product was then prepared for sequencing using the Illumina TruSeq DNA library preparation protocol. Sequencing was performed using the Illumina MiSeq platform, raw sequence files can be found at TBD (Submitting to SRA). Amplification, library preparation and sequencing was performed by Molecular Research LP (Shallowater TX).

### Bioinformatic Pipeline

Raw demultiplexed DNA sequences were first re-orientated and had primers removed using Cutadapt (59). Forward and reverse sequences free of primers and barcodes were trimmed for quality, denoised, merged and grouped into ASVs using the DADA2 pipeline (60). ASVs were assigned taxonomy using the Ribosomal Database Project (27). ASVs that were classified as an unassigned Kingdom or determined to be of Archaeal, mitochondrial, or chloroplastic origin were discarded from the dataset. A phylogenetic tree of ASVs was constructed using the Decipher (61) and Phangorn (62) in R (32). An ASV table, taxonomy table, metadata table, and phylogenetic tree were imported into Pyloseq (28) for further community analysis. Richness estimates were calculated using Breakaway (29) and Shannon estimates were calculated using DivNet (30). Beta diversity was calculated using both weighted and unweighted UniFrac (31), and taxonomic composition relative abundances were computed using Phyloseq. Non metric multidimensional scaling (NMDS) calculations, PERMANOVA analysis of beta diversity distance matrices, CAP, weighted regressions and spearman correlations was computed using the R package Vegan (33). Metabolic inferences were made using PAPRICA (35) by first randomly sampling to an even depth of 9499 sequences per sample. In addition to rarefying sequence counts were normalized using DESeq2 (34) and run through PAPRICA without random subsampling and similar trends were observed.

## ACKNOWLEDGEMENTS

NM gratefully thanks the Science and Resilience Institute at Jamaica Bay for their support of this research, and CUNY for conference travel and dissertation year support. SEA gratefully acknowledges support from NSF-1433014 and PSC-CUNY TRADA-48-463.

Additional support was provided by a Hudson River Fund award (013/15A) from the Hudson River Foundation. We appreciate the support and constructive advice provided by Gateway National Recreation Area. We are thankful for field assistance from Siena Schickler, Dominick Prudente, and Crystal Mena. We also appreciate laboratory and field assistance of the students from the Rockaway Waterfront Alliance Environmentor Program.

---

**Table S1.**
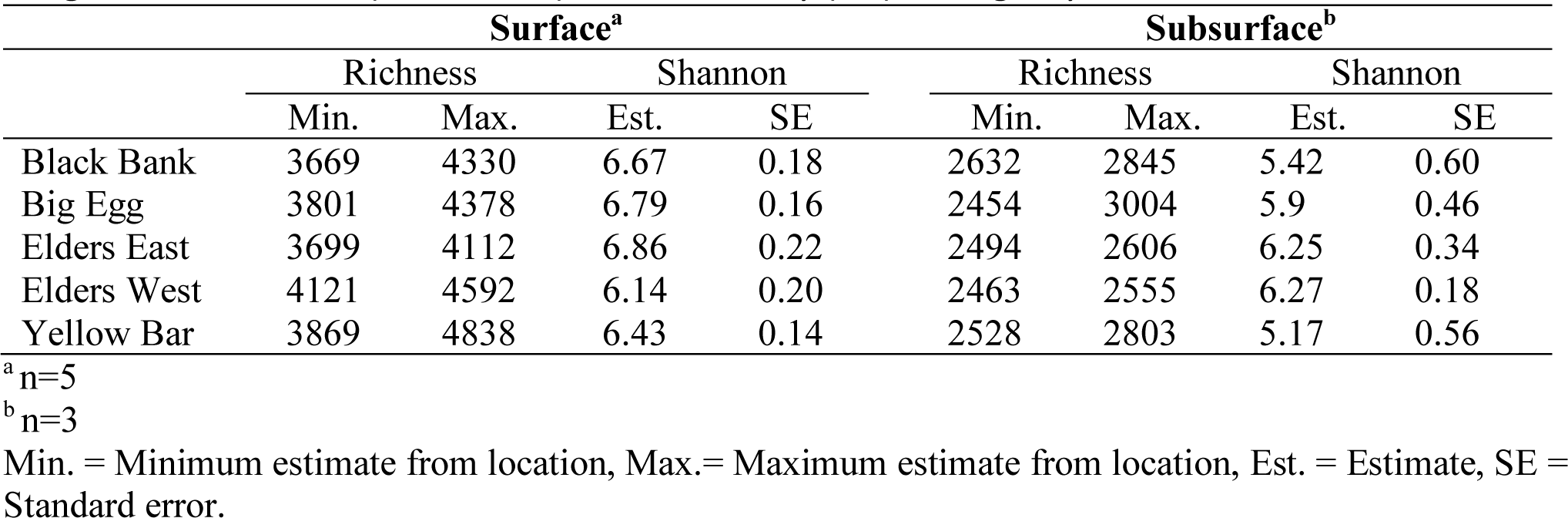
Shannon entropy and richness estimates at four restored salt marshes and a natural degraded salt marsh (Black Bank) in Jamaica Bay (NY) during July 2015….

